# An inexpensive air stream temperature controller and its use to facilitate temperature controlled behavior in living *Drosophila*

**DOI:** 10.1101/439414

**Authors:** Ryan Sangston, Jay Hirsh

## Abstract

Controlling the environment of an organism has many biologically relevant applications. Temperature-dependent inducible biological reagents have proven invaluable for elucidating signaling cascades and dissection of neural circuits. Here we develop a simple and affordable system for rapidly changing temperature in a chamber housing adult *Drosophila* melanogaster. Utilizing flies expressing the temperature inducible channel dTrpA1 in dopaminergic neurons, we show rapid and reproducible changes in locomotor behavior. This device should have wide application to temperature modulated biological reagents.

**Method Summary:** We develop widely applicable and affordable solution to rapidly changing temperature within an enclosed chamber using commercially available components.

## Introduction

Conditional manipulation of effector genes has improved our ability to dissect signaling pathways and neural circuits in living animals. Widely used methods of conditional manipulation include chemical, light, and temperature dependent tools [1]. In *Drosophila melanogaster*, temperature dependent tools such as the *shibire*^*ts*^ allele [2], Gal80^ts^ [3,4], and UAS-dTrpA1 [5] are commonly used. A key benefit afforded by conditional tools is that they allow for temporal control in a constant genetic background. A conditional gene product that is optimal for behavioral studies should be rapidly acting and reversible. Having temporal control affords additional benefits, such as studying genes that may be critical to the viability of an organism during development but disposable later in life.

The original temperature dependent genes relied on naturally existing promoters that responded to heat, for example the heat shock promoter [6]. With the advent of more sophisticated temperature inducible tools, such as heat-activated cation channel dTrpA1, dissecting neural circuitry can be accomplished. In *Drosophila*, the GAL4/UAS binary expression system [4] is widely used to ectopically express these inducible channels in subsets of neurons to measure the physiological and behavioral changes that occur after changing temperature. One of the first demonstrations measured paralysis after inducing dTrpA1 pan-neuronally with the C155-GAL4 driver [5]. Since then, it has been used in many studies defining neuronal circuits such as those involved in learning and memory [7,8], Neuropeptide F signaling [9], and activity levels while free running [10]. Recently a massive screen used 2,204 GAL4 lines to identify neurons responsible for sensory processing, locomotor control, courtship, aggression, and sleep [11]. Despite its widespread usage, methods for inducing dTrpA1 via temperature shifts vary wildly between laboratories and often involve physically moving the animals between environments preset at the different temperatures, which itself can cause behavioral affects due to mechanical stimulation.

In our efforts to measure activity levels of free running flies while conditionally activating dopamine neurons with dTrpA1, we ran in to three issues: first, devices that can change temperature *in situ* are often prohibitively expensive. As with many in the field, we therefore resorted to physically moving animals between chambers set at different temperatures. This led to our second problem, finding confounding phenotypes from mechanical agitation resulting from the movement. Third, published methods for changing temperature are often slow [5,12].

## Results

To address these issues, we developed an inexpensive device that allows rapid temperature changes in a 0.13 m^3^ fly chamber, which is itself housed in a constant environment room (Fig 1). As shown below, temperature shifts are rapid and reproducible, resulting in highly reliable measurements of behavior.

**Fig 1.**
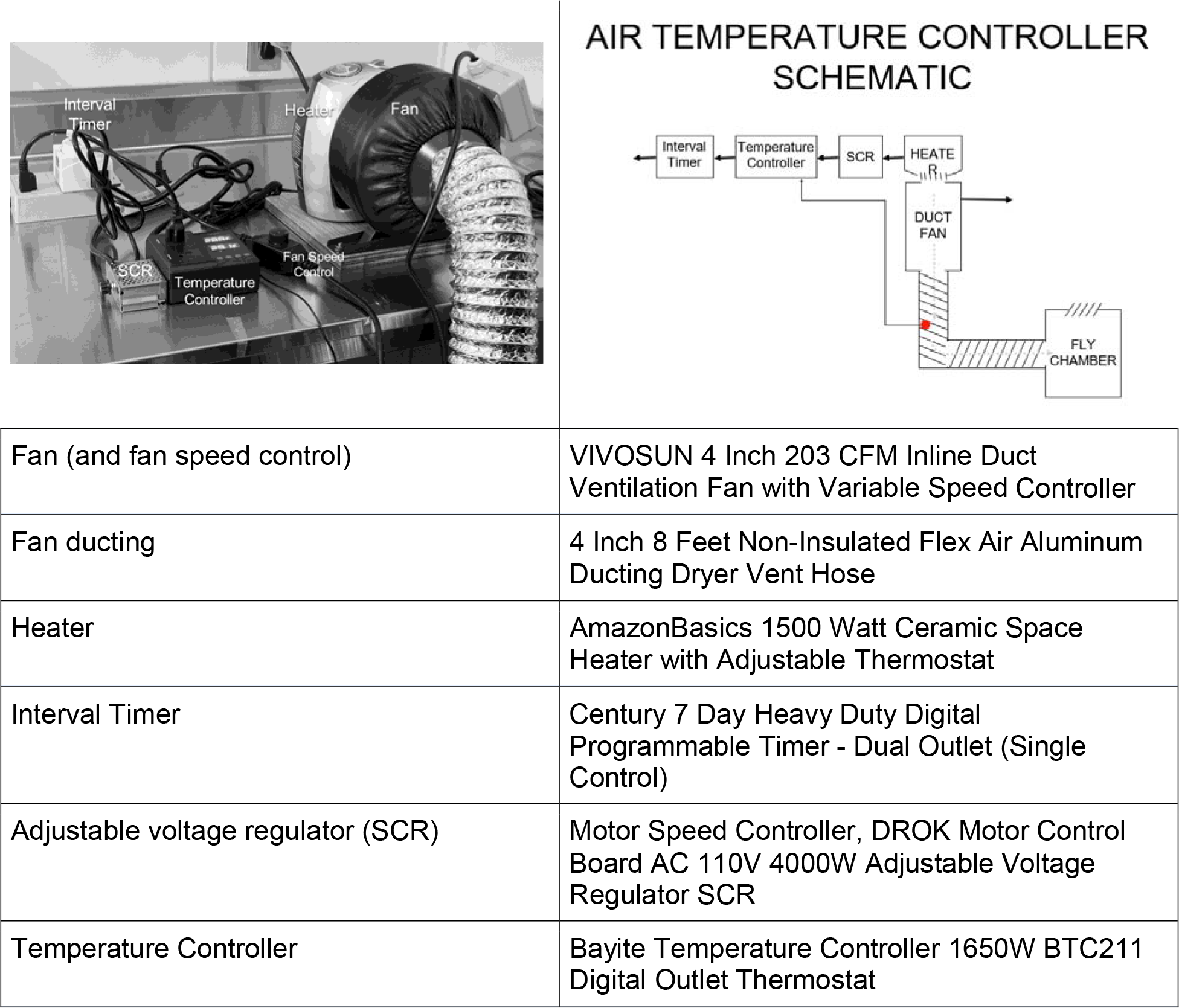
Image and schematic of the air stream temperature controller and parts list. The heat source is an office floor heater, controlled by an interval timer which powers a temperature controller. The power output from the temperature controller is limited by a silicone-controlled rectifier (SCR), to reduce temperature overshoot. The probe (red dot) for the temperature controller is placed within the ducting. The fan is powered continuously and arrows indicate male plugs. The equipment listed is readily available from online retail vendors.

To demonstrate the effectiveness of our system, we measured locomotor responses subsequent to activation of dTrpA1 in most dopamine neurons, using flies containing the TH-GAL4 driver [13] and UAS-dTrpA1 (Fig 2). Dopamine is generally a positive regulator of locomotor activity [14,15]. Therefore, we measured activity upon stimulation using the Drosophila Activity Monitor system (Trikinetics, Waltham, MA). Flies were housed for at least 1 day at 21C, 60% RH, prior to temperature shifts with a 12 hr light/dark schedule, then exposed to 20 min alternating intervals of 29C and 22C in constant darkness but during the subjective daytime. A probe within the chamber indicates that sequential temperature shifts are rapid and highly reproducible (Fig 2), with significant temperature increase detectable by the second minute after activation of the heater, and then stabilizing at an average temperature of 28.7C± 0.6C SEM for the remaining 18 minutes. Temperature decreases are slightly slower due to cooling controlled solely by expulsion of hot air through the chamber vent as opposed to active cooling, but the half time for temperature decrease is still only about 3 minutes. By the fourth minute of the cooling cycle, the temperature falls to under 24C, at which point dTrpA1 is no longer induced. Corresponding activity data from free running flies in response to the temperature shifts is also highly reproducible, with activation of locomotor activity in the UAS-dTrpA1; TH-GAL4 flies coincident with the temperature rise. A one-way ANOVA followed by a Tukey’s honest significance difference post hoc test shows significant increase in activity in driven flies compared to controls from about the second minute at 29C until the tenth minute at which point it decays gradually.

**Fig 2.**
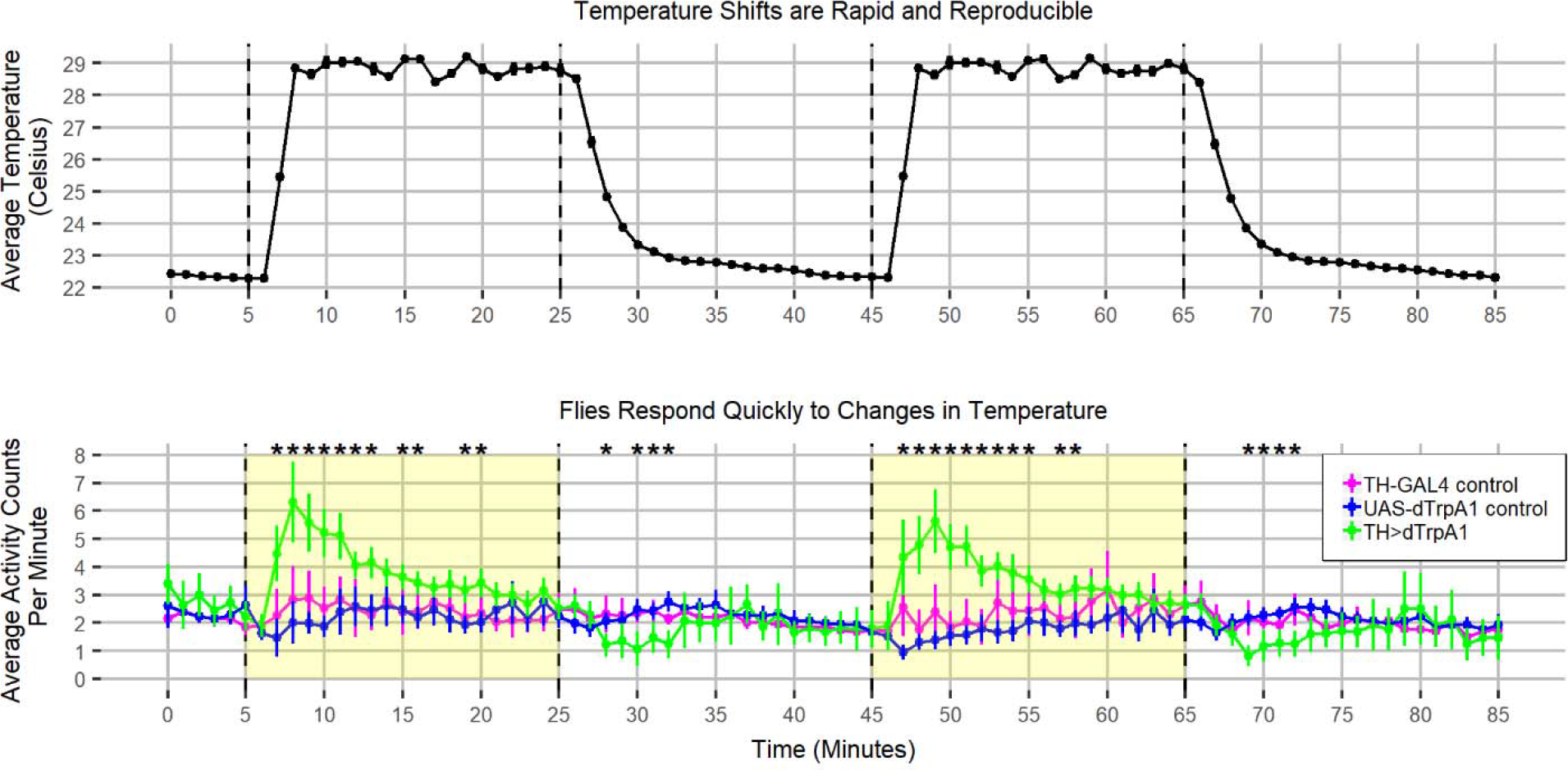
Rapidly induced temperature changes elicit rapid responses from flies expressing dTrpA1 in TH neurons. (Above) Each time point is an average of eight data points at that point in time across eight different days. Dotted vertical lines indicate when the heater was turned on or off by the programmable timer. Error bars are SEM. (Below) Intervals when the heater is activated are highlighted in yellow. By the second minute, TH driven dTrpA1 flies have increased activity levels and the timing coincides with the measured temperature increase. Asterisks indicate a significant difference between driven and control flies where p<0.05 as calculated by a one-way ANOVA followed by Tukey’s honest significance difference post hoc test. TH-GAL4 control n = 31; UAS-dTrpA1 control n = 32; TH>dTrpA1 n = 32.

## Discussion

The locomotor activation resulting from dTrpA1 activation with our paradigm demonstrates its usefulness by uncovering an interesting biological phenomenon: the initial locomotor activation induced by activation of dopaminergic neurons spikes rapidly, then decays well before the 20-minute stimulation is completed. This decay extends into the subsequent downshift to 22C where driven flies exhibit lower activity than the controls. A reasonable explanation for these effects would be that dTrpA1 activation is depleting intracellular stores of dopamine, an explanation that is currently under investigation.

This system also effectively solves three common problems with use of dTrpA1. First, all components are readily available online retail vendors, with a total price ∼$200. Second, it eliminates confounding activity artifacts that result from physically moving flies between chambers. Finally, it improves the time resolution of dTrpA1, with activity responses coincident with the measured temperature rise.

Our system is adaptable to other temperature dependent assays. For example, the digital temperature controller used here has an option for powering a cooling circuit, opening possibilities for more rapid and controlled temperature downshifts. Also, the silicone-controlled rectifier (SCR) can be adjusted to limit the voltage to the heater. This will change the kinetics of the system such that heating occurs quicker or slower based on the biological requirements.

## Experimental Procedures

Parts used are listed in Figure 1.

Flies, in 5 mm OD tubes mounted in DAM2 Activity Monitors (Trikinetics, Waltham, MA) were housed in a custom built 0.13 meter^3^ light-tight wooden container of inner dimensions 54cm W × 54cm D × 45cm H, in a temperature/humidity-controlled walk in environmental chamber (EGC, Chagrin Falls, OH) set at 21C and 60% relative humidity. The EGC chamber was modified to replace the squirrel cage fan coil motors with rotary fans, since the squirrel cage fans generated a strong 120 Hz vibration that affected behavior of the flies. The wooden container has a light tight baffled vent on the side and a front door that opens. It can hold ∼20 fly activity monitors, each capable of monitoring 32 flies. Temperature was monitored in real time using a Temperature Alert monitoring system (No longer sold), or with a Hobo temperature monitor (Onset Computer Corporation, Bourne, MA) set to monitor temperature at 1 min intervals, with the probe placed within the fly chamber.

Temperature conditioned air was ported through the front door of the container via a 4” hole and a clothes dryer vent, connected to 8’ of dryer vent hose. The temperature control unit consisted of an Amazon basics 1500 W ceramic space heater (Amazon), set on its low setting, 750 watts, but with the thermostat set at its high extent. The space heater was mounted adjacent to a duct ventilation fan with double sided outdoor mounting tape, but not physically coupled, since the fan efficiently captures the warmed air.

The temperature was regulated by Bayite temperature controller, set at 29.1C with the heating differential set at 0.1C. The temperature controller output was connected to a voltage regulating SCR set at ∼82V AC, such that the wattage of the heater was ∼510 watts, to prevent overshooting of the setpoint. The SCR output connects directly to the heater. To achieve rapid cycling of temperature, the heater circuit was regulated by a programmable timer, with the fan powered on continuously. Heat generated by the fan motor was sufficient to raise the temperature in the fly chamber ∼1C from the ambient temperature.

Activity data from the Trikinetics monitors was analyzed using a custom R script that averages 5 days of data at each time point and plotted using ggplot2. A one-way ANOVA was calculated at each time point with an alpha of 0.05. At points where a significant difference was detected, a Tukey’s honest significance difference post hoc test was performed. Asterisks indicate time points at which driven flies are significantly different from both controls. Temperature data was also analyzed in R. Eight days of temperature data were averaged at each minute for the indicated temperature shifts.

## Funding Acknowledgements

1. Research reported in this was supported by NIGMS of the National Institutes of Health under grant number R01GM84128.
2. The content is solely the responsibility of the authors and does not necessarily represent the official views of the National Institutes of Health.
3. The authors declare no competing interests.

